# Spatio-temporal learning in a mass-recruiting leaf-cutting ant (*Acromyrmex subterraneus*)

**DOI:** 10.1101/2024.06.12.598721

**Authors:** Fernanda Tiemi Nakashima Ferreira, Pedro Brisola Constantino, Marcelo Arruda Fiuza de Toledo, André Frazão Helene

## Abstract

The ability to anticipate periodically available resources is observed in several animals and improves performance in obtaining resources and adaptability. Spatio-temporal learning occurs when they associate the correct time and location of future events. We evaluate whether leaf-cutting ants *Acromyrmex subterraneus* can anticipate the presence of sucrose and examine potential anticipatory effects. Five colonies were used in an experimental setup where, from a central tray, two trails in opposite directions gave access to either the sucrose or to nothing. For 21 consecutive days sucrose was offered at the same place and the same time. Cameras recorded the flow of individuals at 4 different phases: before feeding (10h00-11h00), pre-feeding (11h30–12h00), feeding (12h00-13h00) and after feeding (14h00-15h00). On the 22^nd^ day sugar was not supplied. On the 21^st^ day the ants were marked and the next day, observed. Our results shows that: (1) the ants responded positively to the stimulus presented by forming the foraging trail to collect sucrose; (2) before feeding there was no significant difference in ant flow between the trails, but after feeding, the ant flow was consistently higher on trail that led to sucrose, which we called a keep going behavior; (3) there was a progressive spatio-temporal learning, given that ants began to appear earlier in pre-feeding throghout the weeks; (4) on the 22^nd^ day, the ants presented themselves 10 minutes in advance and remained in the correct place; (5) the marked ants indicated that even without any resource the empty place continued to be explored. The colonies were able to learn where and when to look for food. Due to the adjustment of the ants to the stimuli of the environment it was possible to prepare for the collection of sucrose and the success in foraging for the colony.

**Summary statement:** Leaf-cutting ants are capable of spatio-temporal learning and this process has relevance on mass foraging recruitment and overall social organization of the colony.

## INTRODUCTION

In nature, food, a crucial resource for any animal, is not available in an unlimited manner and numerous adaptations respond to the challenge of obtaining it. Linnaeus’ floral clock (Gardiner, 1987) demonstrates the adaptive value of anticipating floral opening patterns, crucial for finding resources under specific environmental conditions (Crystal, 2009). In fact, the ability to anticipate the search for periodically available resources is observed in several animals, including rats (Wall et al., 2019), birds (Tello-Ramos et al., 2015), bees (Murphy and Breed, 2008; SILVA et al. , 2008; SILVA et al., 2021) and neotropical ants (Schatz et al., 1999). Anticipation improves performance in environmental challenges (Reznikova, 2007), allowing preparation for stimuli and increasing the chance of a successful response (Cipolla-Neto et al., 1988).

Spatio-temporal learning, assessed experimentally by training animals to feed based on defined spatio-temporal contingencies, involves associating the correct timing and location of future events (Beugnon et al., 1995; Fourcassié et al., 1999). This anticipatory activity targets objects or places that provide food (Mistlberger, 2009). The benefits of this learning process are profound, deeply rooted within the endogenous circadian rhythm system found in all animals. This system synchronizes with environmental cycles, responding to periodic stimuli that align with day/night patterns (Aschoff, 1989; Beugnon et al., 1995). Such synchronization enhances the adaptive capabilities of animals, illustrating the intricate interplay between internal biological clocks and the external world.

In fact, experiments with eusocial animals such as bees (Fourcassié et al., 1999) and *Ectatommaruidum* ants (Schatz et al., 1999) show that they can associate feeding sites with different times, with ants beginning to visit the same ones locations at different times, up to 30 minutes before food was made available. It is reasonable to assume that this characteristic is associated with ecological factors that affect the natural availability of food sources, explaining why neotropical ants can learn when and where food is available (Harrison and Breed, 1987).

However, in neotropical leafcutter ants, mass collective foraging adds a dimension of sociability to spatio-temporal learning, allowing the encoding of information about food sources on the collectiveness.

In this context, the present work aims to evaluate whether ant colonies *Acromyrmex subterraneus* can anticipate the presence of sucrose and examine potential anticipatory effects related to individual and mass foraging behaviors.

## MATERIALS AND METHODS

For the study, 5 colonies of *Acromyrmex subterraneus* of at least one-year old, bearing 1200 to 1800 ml fungus garden with 1.000 to 1.250 ants by colony, were used. They were kept in the laboratory under controlled conditions of temperature (25°C), relative humidity (70%) and photoperiod 12:12 LD. Each colony was maintained for acclimatization for 5 days in the experimental setup (described below), when the ants could freely explore the new environment.

### EXPERIMENTAL SETUP

The experimental arena consisted of three trays connected through wooden bridges. The main colony, containing the fungus garden, was placed in the central tray. Sucrose solution was offered at a fixed phase (feeding phase) in one of the other peripheral tray (choose randomly each trial), which we will refer as sucrose tray hereinafter. The other peripheral tray was kept empty, which we will refer as empty tray hereinafter. Two recording webcams (Logitech c920 pro full HD) were placed at the end of both bridges connecting the trays. The bridge connected to the empty tray was called the Empty Trail and the other connected to the tray where the food was placed, the Sucrose Trail. These webcams were connected to a computer running an open license surveillance software (iSpy®) in order to record the ant flow at the bridge during the experiment selected phases (figure 1).

**Figure 1:**
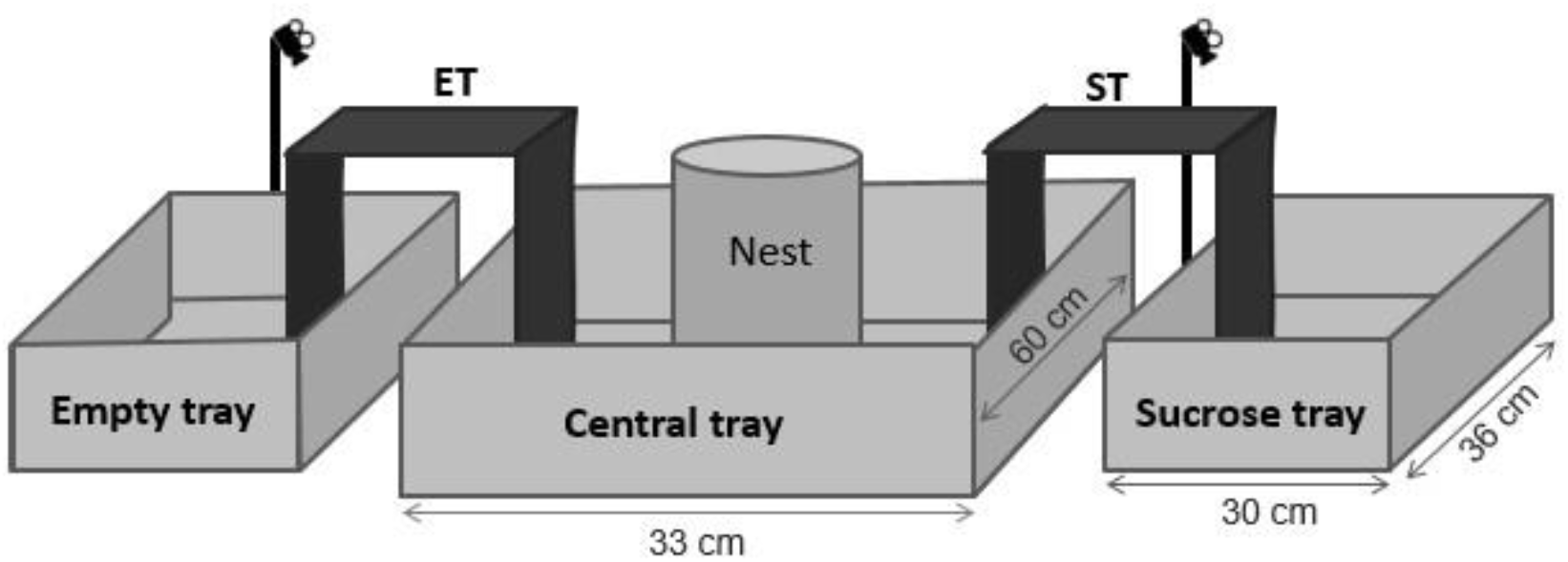
Representation of the experimental setup. Central tray containing the nest connected by two bridges: empty trail (ET) that leads to the empty tray and sucrose trail (ST) that leads to the sucrose tray.

### EXPERIMENTAL PROCEDURE

During the entire experiment, ants were fed *ad libitum* with *Acalipha* sp. leaves and oat flakes in the central tray. At this stage, the iSpy image recording program was used, programmed for daily filming of videos organized into 10-minutes videos at the pre-established phases at which the filming took place: before feeding (10h00 to 11h00); pre-feeding (11h30 to 12h00); feeding (12h00 to 13h00), when sucrose solution was available at sucrose tray; and after feeding (14h00 to 15h00).

The experiment lasted 22 consecutive days. For 21 consecutive days, approximately 1.5 ml of 0.6M sucrose was placed on a clean glass plate (8 cm x 4 cm) at 12h00 in the sucrose tray. When removing the sucrose at 13h00, another clean plate was placed in its place. On the 22^nd^ day, sucrose was not provided.

### ANT FLOW COUNTING

The ant flow on the recorded videos was analyzed using automatic animal flow counting software developed in the laboratory.

To analyze the videos, a movement filter is first applied, which separates the pixels in which there was movement (foreground) from the static ones (background) using the ViBe algorithm (Barnich and Van Droogenbroeckref, 2011). Then, the centroids of the connected components are obtained. After this step, to establish the correspondence between which centroids in a frame correspond to the centroids of the following frame (tracking), the R-RANSAC algorithm was used (Niedfeldt and Beard, 2014). Finally, on the filmed frame, a line was positioned, for each trajectory that crossed this line, but did not cross it back; a count was recorded for the respective direction. The implementation of these analyzes was done in C++, using the OpenCV library.

### ANT MARKING

To observe the individual behavior of the ants, the individuals present in each tray at the 21^st^ day were marked with an oil-based sharpie pen following a unique pattern, represented in table 1. The ant marking period involved only the “pre-feeding” and “feeding” phases. On the 22^nd^ day, ants marked during “pre-feeding” and “feeding” were manually counted in the empty and sucrose trays every 15 minutes.

**Table 1:**
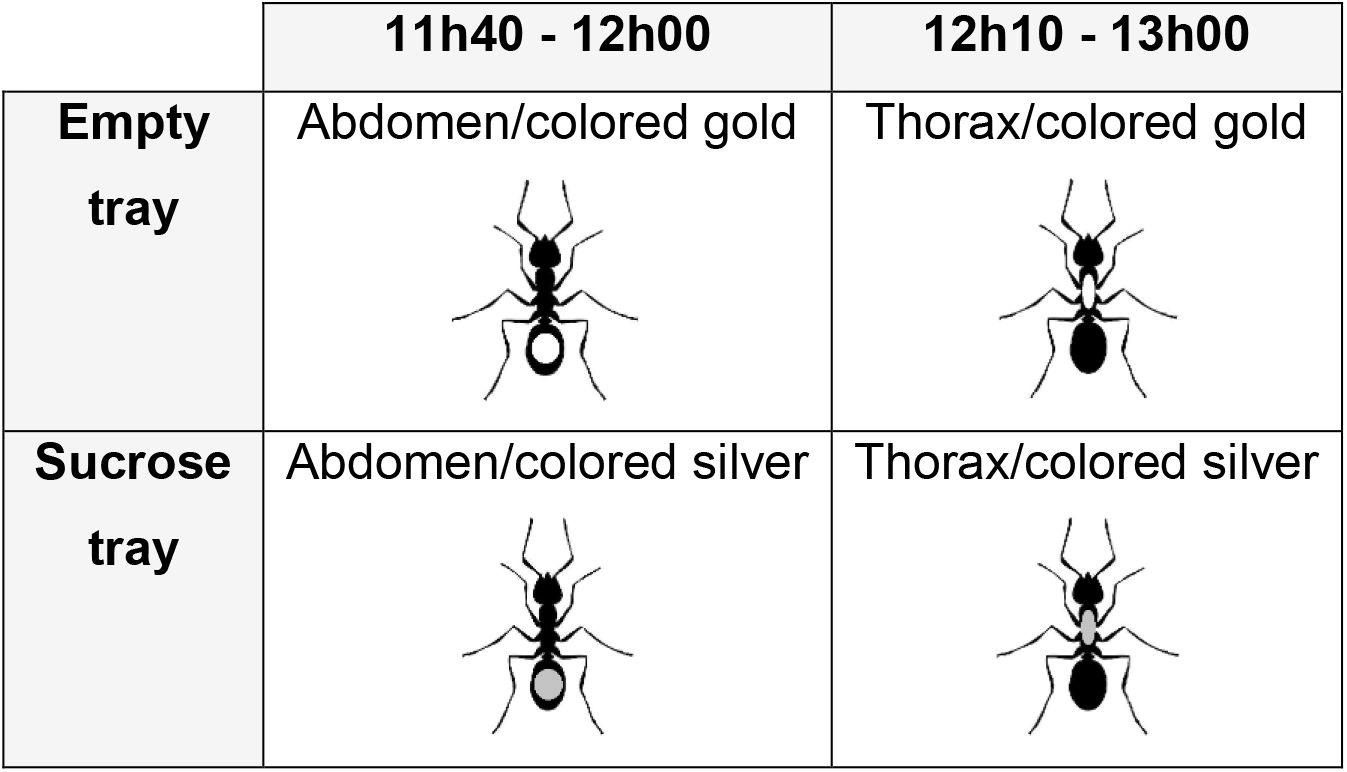
Ant marking scheme in foraging trays.

|  | 11h40 - 12h00 | 12h10 - 13h00 |
| --- | --- | --- |
| Empty tray | Abdomen/colored gold | Thorax/colored gold |
| Sucrose tray | Abdomen/colored silver | Thorax/colored silver |

### DATA ANALYSIS

Statistical analysis was performed using the Friedman Test with the RStudio graphical interface package version 1.2.1335. The graphics also were created by RStudio.

The Friedman test was performed with the average colony flow and with time intervals of 10 minutes. Finally, the difference in ant flow between foraging trails, between experimental days, between weeks, between established times and between minutes were compared.

To analyze individual ant behavior on the 22^nd^ day we counted total number of marked ants on each foraging trail at pre-feeding and feeding phases. For each group of marked ants, binomial distribution analysis was performed, with a significance of p<0.05 both at the time referring to “pre-feeding” and “feeding”.

## RESULTS

### OVERVIEW OF ANT FLOW

The offering of sucrose caused general changes in the ant flow. In Figure 2A and 2B, it is possible to observe the average value of ant flow for all colonies over the 21 days, at each time and in both trails. The most striking feature of our results is that sucrose availability increased flow on sucrose trail while decreased on empty trail.

**Figure 2:**
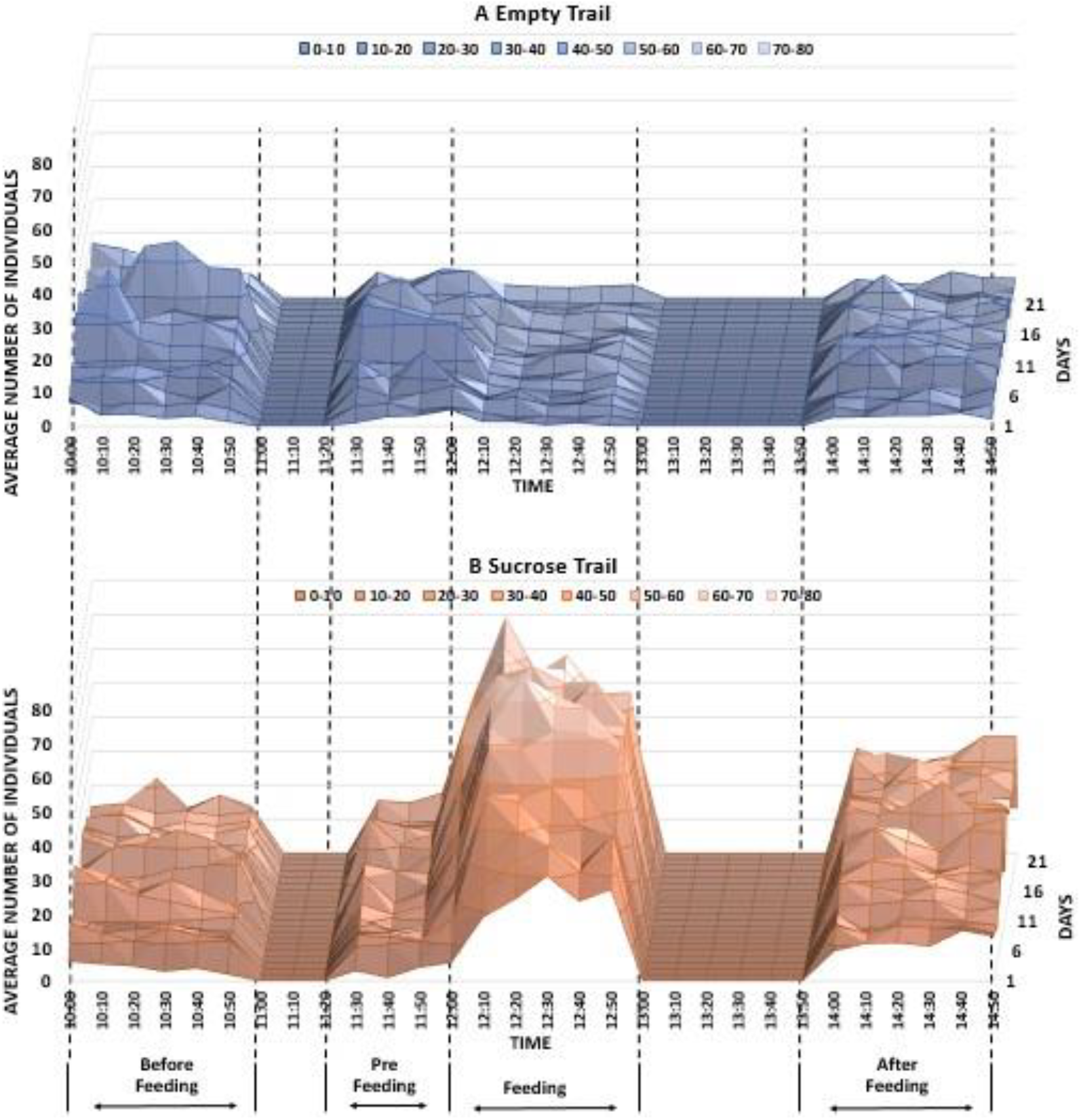
Ant flow in the 22 days of the experiment at times “before feeding”, “pre-feeding”, “feeding” and “after feeding” on the (A) EmptyTrail and (B) Sucrose Trail.

**Figure 3:**
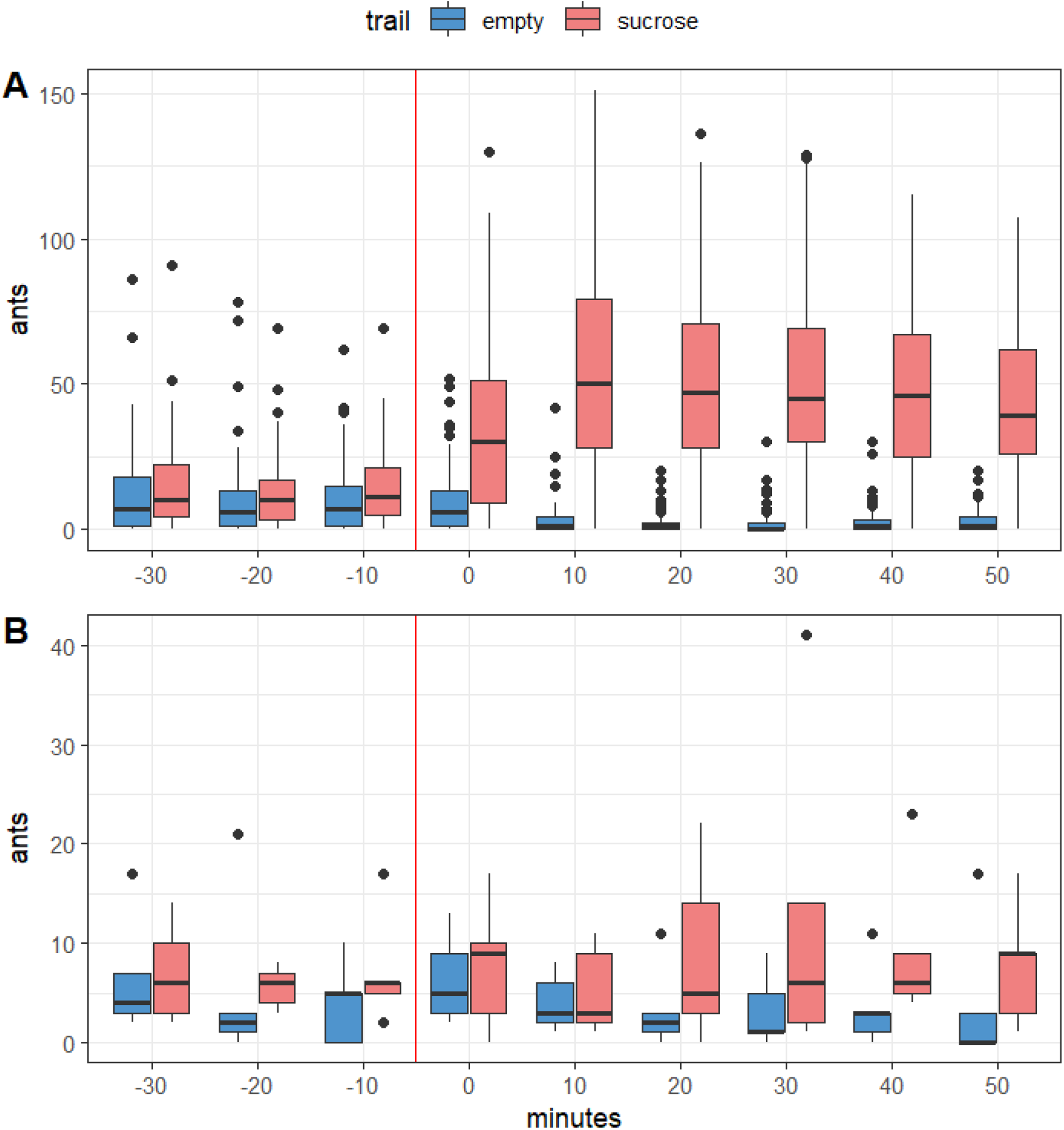
number of ants counted in sucrose (coral) and empty (blue) trail throughout the 21 days of experiment (A) and on the 22^nd^ day (B). Red vertical line represents the instant food was offered.

### FORAGE ON THE TRAILS

As cited above, sucrose changed ant flow in both trails, increasing it on the sucrose trail, while decreasing it on empty trail. A deeper analys shows the effect of spatio-temporal learning being built over the 21 experiment days, as ant flow to sucrose trail began to increase 10 minutes before sucrose was made available by the end of the first week (see bellow for more information). Moreover, on the 22^nd^ day the effect persisted, showing that indeed there were few ants anticipating sucrose offering.

The weekly analysis allows us to verify that, during the “pre-feeding”, the difference between the flow of individuals on the empty trail and sucrose trail varied. In the first week, this difference was not observed. In the second week there was a significant difference at 11h50 and in the third week it happened at 11h40 and 11h50 (Figure 4). These data indicate a progressive change in behavior signaling a construction of anticipation.

**Figure 4:**
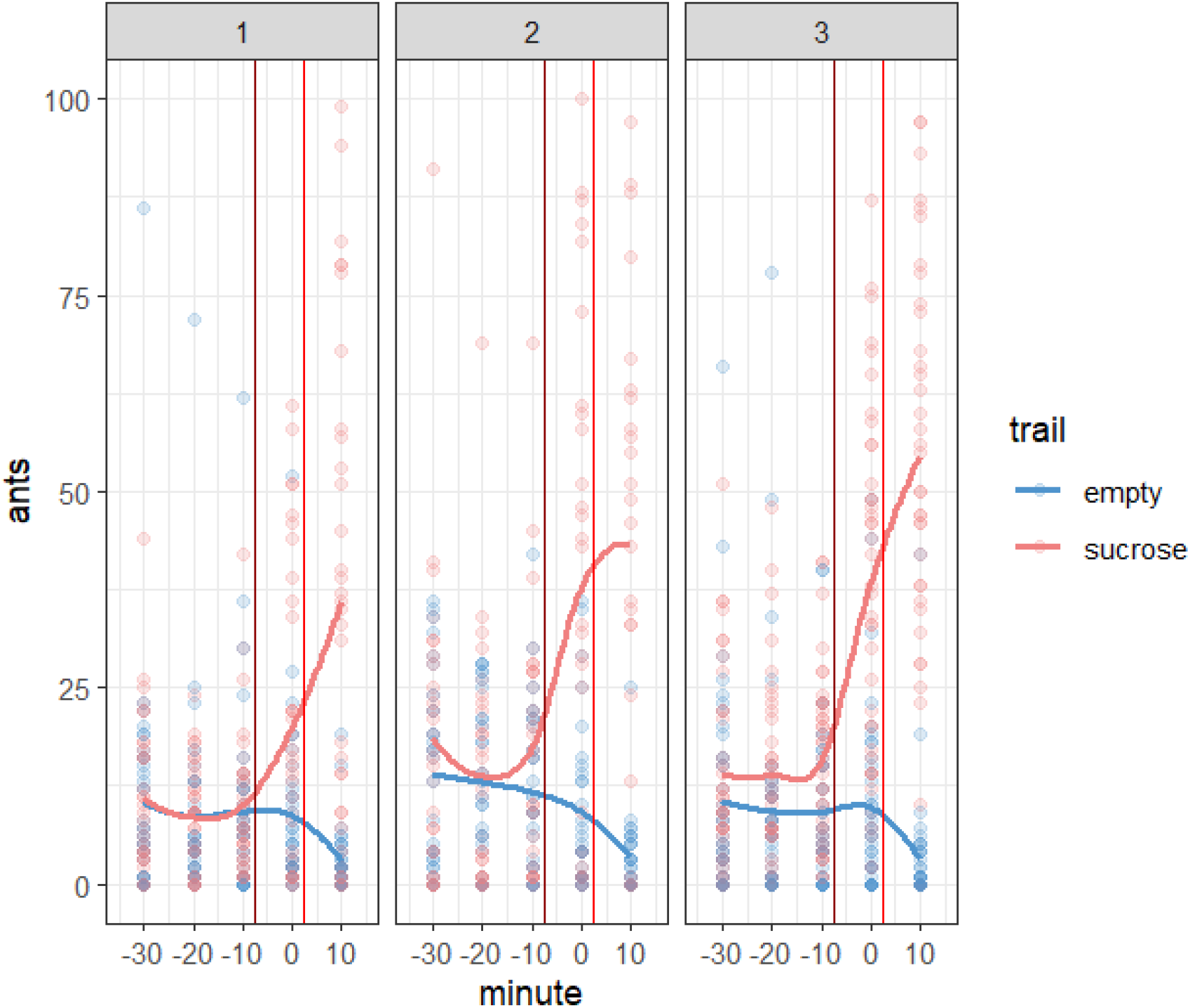
number of ants counted at trails leading to both sucrose and empty feeding trays during pre-feeding phase (first 10 minutes of feeding phase also showed). Data sampled from day 1 to 21 but divided in 3 weeks (panels 1, 2 and 3). Coral and blue lines represents a local regression (LOESS locally estimated scatterplot smoothing) fitted minute-to-minute to the data. Red vertical line represents food offering and burgundy red line indicates possible anticipation inflection point (see results and discussion for further analysis).

In before feeding phase there was no difference in the ant flow between the foraging trails (Figure 5A). After feeding, a latent post-sucrose flow was observed on the sucrose trail, causing a significant difference between the foraging trails on the days of the experiment in which sucrose was added. The same pattern was observed on the 22^nd^ day before feeding (Figure 5B). However, as sucrose was not supplied on this day, the effect was significantly smaller than when there was sucrose.

**Figure 5:**
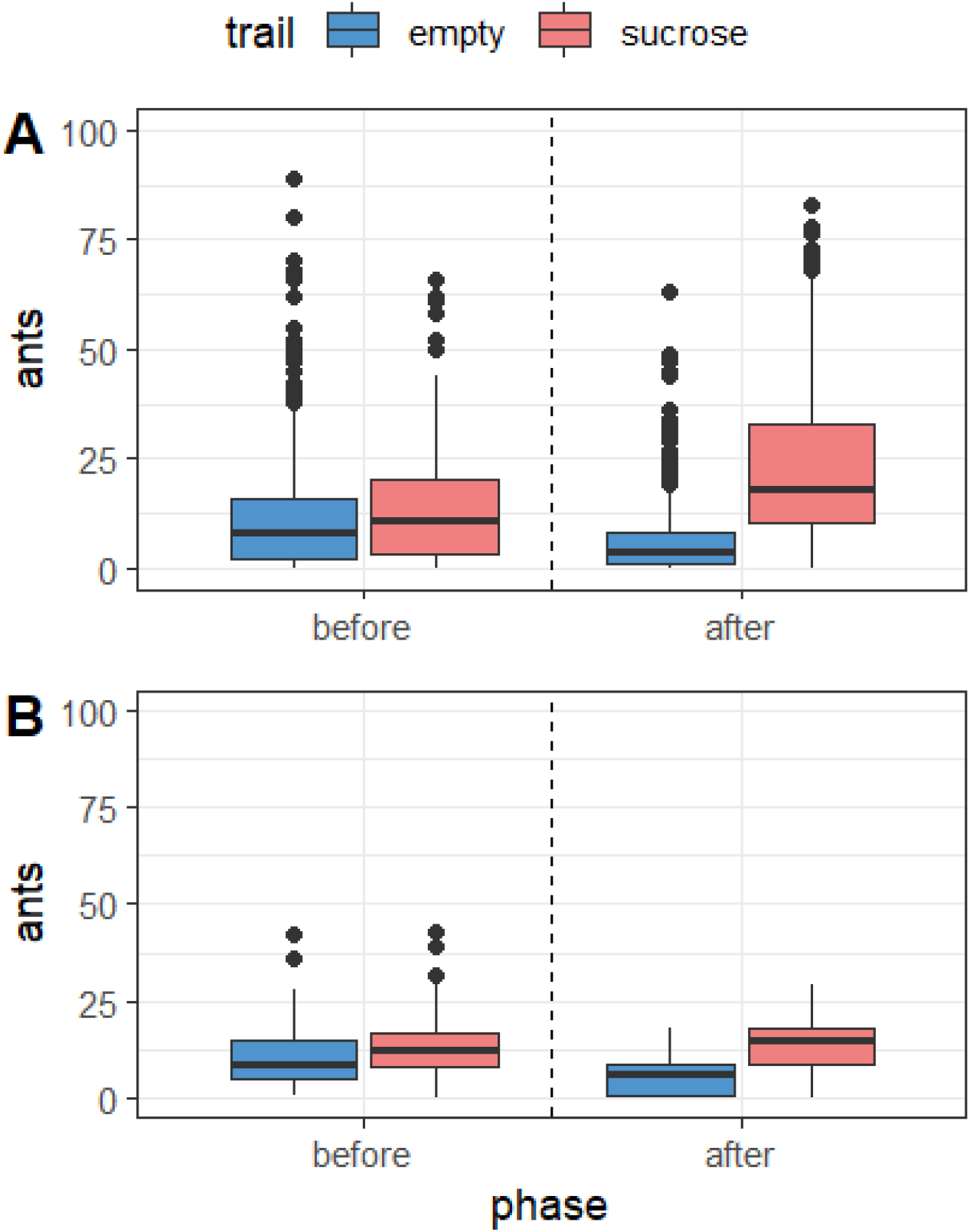
Number of ants counted on sucrose (coral) and empty (blue) trails before and after feeding throughout the 21 days of experiment (A) and on the 22nd day (B).

### ANALYSIS OF MARKED ANTS

A total of 427 ants were marked on the 21^st^ day (Table 2). They were observed on the 22^nd^ day and separated into four groups according to the marking pattern that can be seen in Table 1.

**Table 2:** Number of ants marked in the foraging trays.

| Appointment<br>time | Number of ants marked |  |
| --- | --- | --- |
|  | Pre-feeding<br>11h40 - 12h00 | Feeding<br>12h10 - 13h00 |
| Empty tray | 50 | 40 |
| Sucrose tray | 115 | 222 |

In Figure 6, the data presents the four groups that were observed in the trays during pre-feeding and feeding. The ants which was marked in the empty tray from 11h40 to 12h00, showed a significant difference in the number of ants during “pre-feeding” and “feeding”, with a preference for this tray (binomial test with expected probability of success of 0.50: “pre-feeding”: 12/17, Z= 9.62, p=0.047; “feeding”: 22/33, Z=7.66, p=0.022).

**Figure 6:**
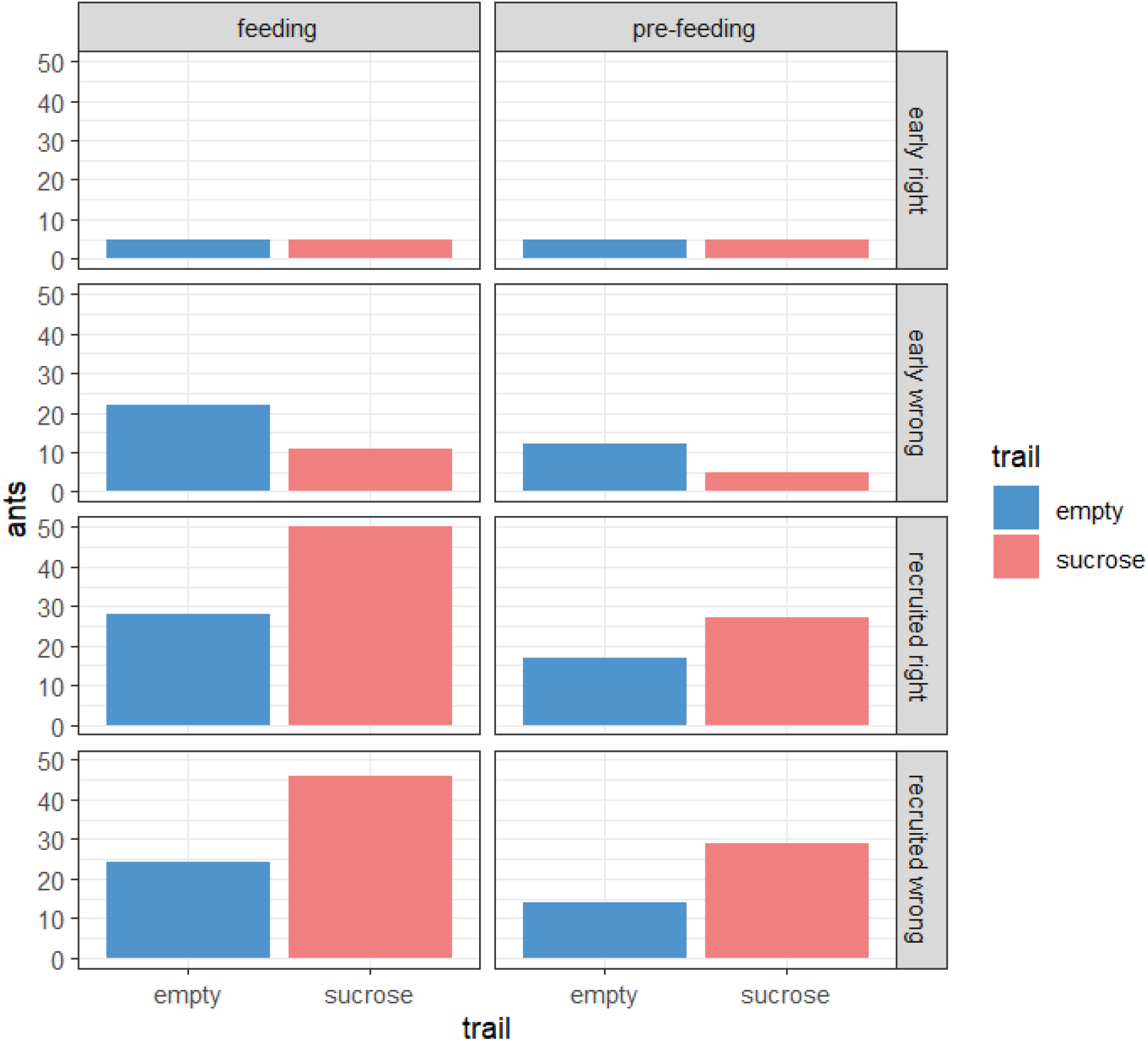
Number of ants observed in the trays on the 22^nd^ day of the experiment.

The individuals marked on the sucrose tray were observed in greater numbers on that same tray, also at both times when data collection was carried out (group from 11h40-12h00: “pre-feeding”: 24/38, Z=5.98, p=0.035; “feeding”: 46/75, Z=5.13, p=0.013; group from 12h10-13h00: “pre-feeding”: 27/44, Z=5.14, p<0.04; “feeding”: 50/78, Z =6.43, p<0.005).

## DISCUSSION

The experiments showed that the presence of sucrose changed the colony behavior in many different ways. First, it caused the number of ants to increase considerably on the Sucrose trail during feeding phase while decreasing on the empty trail. Second, sucrose availability made ant distribution change between the before feeding and after feeding phases: before feeding there was no significant difference in ant flow between empty trail and sucrose trail, but after feeding, the ant flow on sucrose trail was consistently higher than on empty trail. Third, we were able to measure an anticipation behavior building throughout the experiment weeks, as ant flow going to the sucrose tray, measured in pre-feeding phase, began to increase before sucrose offering.

### RESPONSE TO STIMULUS AND ANTICIPATORY BEHAVIOR

When comparing the flow of ants between the foraging trails in the pre-feeding phase, a significant preference for sucrose trail before feeding was verified throughout the 21 days. And the ants learning occurred gradually, with an anticipation of 10 minutes in the second week and 20 minutes in the third week (Figure 4). In rodents, 2 or 3 weeks of experimentation is recommended if anticipation does not occur in the first week (Mistlberger, 2009).

The preference and gradual increase in flow on the sucrose trail shows that the ants adapted to the location where sugar was available. They adjusted their activity level due to environmental stimulation (Gordon, 1996). As leaf-cutting ants do mass foraging, the mechanisms to organize the collective behavior in order to learn spatio-temporal cues are imbedded in the main means of navigation, the integration of paths and reference points (Wehner et al., 1990; Collett, 1992) and our experimental setup was designed for ants to use them.

The increase in “pre-feeding” flow in the sucrose trail can be interpreted as anticipation. Only a few ants may have learned the task by synchronizing their activities with the feeding stimuli, while some other ants may have learned from ants that had already learned (Fourcassié et al., 1999), or both. The dynamics of the colony influence the behavior of the individual (Feinerman and Korman, 2017; Frank and Linsenmair, 2017) due to the patterns of interactions formed during the days in which the stimulus was presented (Gordon, 2018), which justifies gradual spatio-temporal learning. This learning depends on the species and type of food available (Cammaerts, 2013).

When verifying the effect of sucrose on sucrose trail, it is possible to observe that there was a recruitment process during feeding phase, as verified by the high flow of individuals (Figure 3A). Recruitment is common in poneroids, Formicinae, and Myrmicinae ants and honeydew is collected by ants on trails that can persist for months or years and that requires multiple ants to care for the workers engaged in foraging and defend the patch (Lanan, 2014). Therefore, recruitment involves the positive feedback process that causes amplification in the number of ants (Camazine et al., 2001). This recreuitment process was not observed with *Ectatommaruidum* ants, even though they are also capable of anticipating (Schatz et al., 1999).

One hour after the food was removed, it was still possible to observe a considerable flow in the sucrose trail during after feeding phase along the 21 days (Figure 5A), we call this a “keep going” effect. At this stage, the negative feedback process allows the system to “stabilize” again due to food exhaustion (Bonabeau et al., 1997). This explains why the flow is still relatively high after feeding as it is a product of mass recruitment generated by the presence of the stimulus, while before feeding, even with the marked trail; there is no difference in ant flow between the trails. We believe that night plays an important role in resetting the system. In a condition where food was available continuously, the trail would persist nonstop; however, ants passing by it wouldn’t be the same, as they are capable of divide labor in work shifts (Constantino et al., 2022). So, it is possible that, as workers change during night, and they don’t find the food, the collective organization maintaining them on the sucrose trail dismantle, resetting the system. So, during pre-feeding, ants that found food previously, return anticipating the availability of sucrose.

The 22^nd^ day, a day without sucrose, also indicates that there was spatio-temporal learning of the task. Temporal analysis shows that the ant flow was greater in sucrose trail at: “pre-feeding”, 10 minutes before food offering (Figure 3B); in “feeding” the ant flow was greater in all 10-minute intervals, but not as near as when food was available. This shows that the increase in ant flow during feeding phase had two influences: the anticipation behavior, given by a subset of workers; and the recruitment, a significantly greater effect.

On the 22^nd^ day there wasn’t a clear “keep going” effect (Figure 5B), mostly because there wasn’t a strong recruitment, as there wasn’t food offering. The option of choosing sucrose trail, and not empty trail, signals that they were waiting for sucrose at the expected location and time. In other words, there was a preferred trail but there was no recruitment.

The fact that some ants anticipated and decided to stay in the place even without food shows that they sacrifice their time and energy for the good of the colony, as was verified by Roces and Núñez (1993). According to Schatz et al. (1999), the complete or partial absence of reward is associated with a significant increase in travel time, suggesting that ants prefer to remain in the correct location even without the reward rather than moving to another location. Furthermore, it has been seen that there are bees that have learned to anticipate and make periodic visits before and after food is available to spend less energy searching for new food sources (Jesus et al., 2014).

Animals use memory to survive and improve their lives (Reznikova, 2007). In nature, anticipation indicates prevention against food competitors (Schatz et al., 1999; Roces, 2002) greater chances of preparation (Cipolla-Neto et al., 1988), agility in recruitment (Roces, 2002) and greater guarantee of colony survival (Reznikova, 2007).

### CONSIDERATIONS OF SPATIO-TEMPORAL LEARNING IN THE COLONY

It was found that the number of ants that anticipated the presentation of sucrose was low, not involving mass recruitment. As scouts control the behavior of the colony (Frank and Linsenmair, 2017), this activity is possibly related to them. While the other castes are carrying out their activities in or near the nest, the scouts leave the nest to forage and fulfill their role, indicating that anticipation is an individual learning and does not depend on mass recruitment. The division of labor combined with cooperation is a great advantage for social insects (Wilson, 1980) and the increase in the degree of specialization of workers contributes to the organization of the colony (Wilson, 2000). Through some individuals, the memory and learning of ants can be expressed (Morley, 1958).

Furthermore, collecting sucrose consumes time and energy (Carmo, 2014), meaning there is no need for a large number of ants waiting for the sucrose to be placed in the tray. This may explain the low flow on the sucrose trail on the 22^nd^ day (Figure 3B). In *Melipona subnitida* bees there are foragers that do not anticipate avoiding unnecessary predation risks, exposure to high temperatures and low humidity (Silva et al., 2021). So, maybe, there is a trade-off between anticipating and gaining advantage on retrieving resources; and leaving the nest too early and face harsh foraging conditions and predators.

It was also found that *Acromyrmex* colonies responded positively to the time of day in which the food was provided. Although greater nocturnal foraging activity has been observed in *Atta cephalotes* (Cherrett, 1968) and *Atta sexdens rubropilosa* (Toledo, 2013; Constantino, 2017), which are also leaf-cutter ants, it is known that ants adapt to foraging at night in summer due to thermoregulation, and in winter they adjust to leave the nest during the day (Holldobler and Wilson, 2011).

Thus, it was possible to observe that sucrose acted as a *zeitgeber* allowing the ants to synchronize their activities to cyclical events in the environment (Aschoff, 1989). A possible entrainment may have occurred to ensure the colonies foraging success during “feeding”.

### INDIVIDUAL STANDARDS

It was found that ants forage along stereotypical paths when moving between two known places (Collett et al., 1992). The ant marking test corroborated Collett et al. (1992) by showing that individuals that were marked in the empty tray and the sucrose tray were seen in their corresponding trays during “pre-feeding” and “feeding”. With the exception of the ants that were marked in the empty tray in “feeding” (Figure 6). Even though the colonies activities were focused on the sucrose tray due to the stimulus presented for 21 days, the exploration of the other tray was not completely abandoned. There were workers responsible for this activity, as the same ants that were marked in the empty tray were also seen there, preferentially. Probably to explore more resources elsewhere, be aware of competition (Gordon, 2002).

## CONCLUSION

The data collected in this study showed that ants *Acromyrmex subterraneus* are able to progressively anticipate the presentation of sucrose. Behavioral plasticity was important so that the ants’ 20-minute anticipation could happen. This learning occurred individually and did not involve a large number of ants, which possibly prevented the loss of time and energy for the colony as a whole on the day that sucrose was not presented. And even with the focus of foraging being on the sucrose tray, the empty tray continued to be explored, including when sucrose was collected. If each individual plays their role by adapting to the challenges of the environment for the success of the colony, a true society can be observed. The absence of mass anticipatory recruitment shows a separated individual and collective behavior.

## ACKNOWLEDGEMENTS

We would like to thank Professor Odair da Costa Bueno and Center of Studies on Social insects (CEIS-UNESP-Rio Claro) for providing the colonies used in our experiments

